# Repetitive transcranial magnetic stimulation (rTMS) triggers dose-dependent homeostatic rewiring in recurrent neuronal networks

**DOI:** 10.1101/2023.03.20.533396

**Authors:** Swathi Anil, Han Lu, Stefan Rotter, Andreas Vlachos

**Author notes:** Correspondence to (SR), (AV). joint senior authors (Stefan Rotter, Andreas Vlachos).

## Abstract

Repetitive transcranial magnetic stimulation (rTMS) is a non-invasive brain stimulation technique used to induce neuronal plasticity in healthy individuals and patients. Designing effective and reproducible rTMS protocols poses a major challenge in the field as the underlying biomechanisms remain elusive. Current clinical protocol designs are often based on studies reporting rTMS-induced long-term potentiation or depression of synaptic transmission. Herein, we employed computational modeling to explore the effects of rTMS on long-term structural plasticity and changes in network connectivity. We simulated a recurrent neuronal network with homeostatic structural plasticity between excitatory neurons, and demonstrated that this mechanism was sensitive to specific parameters of the stimulation protocol (i.e., frequency, intensity, and duration of stimulation). The feedback-inhibition initiated by network stimulation influenced the net stimulation outcome and hindered the rTMS-induced homeostatic structural plasticity, highlighting the role of inhibitory networks. These findings suggest a novel mechanism for the lasting effects of rTMS, i.e., rTMS-induced homeostatic structural plasticity, and highlight the importance of network inhibition in careful protocol design, standardization, and optimization of stimulation.

**Author summary:** The cellular and molecular mechanisms of clinically employed repetitive transcranial magnetic stimulation (rTMS) protocols remain not well understood. However, it is clear that stimulation outcomes depend heavily on protocol designs. Current protocol designs are mainly based on experimental studies that explored functional synaptic plasticity, such as long-term potentiation of excitatory neurotransmission. Using a computational approach, we sought to address the dose-dependent effects of rTMS on the structural remodeling of stimulated and non-stimulated connected networks. Our results suggest a new mechanism of action—activity-dependent homeostatic structural remodeling—through which rTMS may assert its lasting effects on neuronal networks.

We showed that the effect of rTMS on structural plasticity critically depends on stimulation intensity, frequency, and duration and that recurrent inhibition can affect the outcome of rTMS-induced homeostatic structural plasticity. These findings emphasize the use of computational approaches for an optimized rTMS protocol design, which may support the development of more effective rTMS-based therapies.

## Introduction

Repetitive transcranial magnetic stimulation (rTMS) is a non-invasive brain stimulation method used in basic and clinical neuroscience(1,2,3). Based on the principle of electromagnetic induction, rTMS induces electric fields that activate cortical neurons and modulate cortical excitability beyond the stimulation period (4,5,6). This makes rTMS a suitable tool for studying and modulating brain plasticity in healthy and disease states [7,8, 9, 10,11].

Experiments in animal models have shown that rTMS induces specific changes in excitatory synapses, that are consistent with a long-term potentiation (LTP) of neurotransmission (12, 13, 14, 15). Using animal models (both *in vitro* and *in vivo*), we also previously demonstrated rTMS-induced changes in inhibitory neurotransmission, wherein a reduction in dendritic but not somatic inhibition was observed (16). These findings provide an explanation of how rTMS may assert its effects—by mediating disinhibition and priming stimulated networks for the expression of physiological context-specific plasticity (17). Nevertheless, it remains unknown how exogenous electric brain stimulation that is not linked with specific environmental or endogenous signals asserts therapeutic effects in patients.

In recent years, a considerable degree of variability (or even absence) of rTMS induced “LTP-like” plasticity—measured as a change in the evoked potential of the target muscle upon stimulation of the motor cortex (18, 19, 20, 21)— has been reported in human participants, often leading to difficulties in reproducing results (22). Efforts to explain this variability have largely focused on the assessment of possible confounding factors that may affect the outcome of a given rTMS protocol as well as on prospective optimization of induced electrical fields for standardization of stimulation protocols and dosing across participants (23, 24). This has also led to discussions on alternative underlying mechanisms, such as the impact of rTMS on glial cells and rTMS-induced structural remodeling of neuronal networks (25, 26, 27, 28). There has been emerging evidence of structural plasticity induced by rTMS. Studies have demonstrated that rTMS facilitates reorganization of abnormal cortical circuits (10, 11), which may be pertinent to its therapeutic effects and cognitive benefits (29,30). Moreover, structural connectivity changes induced by rTMS have been shown to underlie anti-depressant effects in chronic treatment-resistant depression (31, 32, 33). Vlachos et al. (12) also demonstrated structural remodeling imposed by 10 Hz repetitive magnetic stimulation on small dendritic spines in an *in vitro* setting. More recently, structural synaptic plasticity in response to low-intensity rTMS was demonstrated using longitudinal two-photon microscopy in the motor cortex of mice (14). Towards this direction, we used network simulations to evaluate the dose-dependent effects of rTMS on the structural remodeling of neuronal networks in this study. We evaluated rTMS-induced structural changes that may occur even in the absence of changes in synaptic weights (i.e., LTP-like plasticity). Specifically, we employed an inhibition-dominated recurrent neuronal network with homeostatic structural plasticity that follows a negative feedback rule (34,35, 36). In this network, continuous synaptic remodeling takes place in order to maintain neuronal activity at a stable point. Deviation from this level of activity are restored using synaptic formation or deletion at regular intervals. Based on our previous experimental findings that 10 Hz stimulation induces structural remodeling of excitatory synapses and dendritic spines (12), we assessed the effects of stimulation intensity, pulse number, and frequency—including clinically established intermittent theta burst stimulation (iTBS)—on rTMS-induced homeostatic structural plasticity.

## Materials and methods

### Neuron model

All large-scale simulations in the present study were performed using NEST simulator 2.20.0 (37), using MPI based parallel computation. Single neurons were modeled as linear current based leaky integrate and fire (LIF) point neurons, having subthreshold dynamics expressed by the following ordinary differential equation:

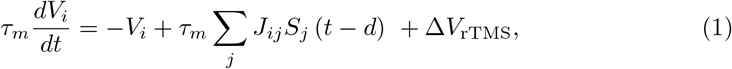

where *τ*_*m*_ is the membrane time constant. The membrane potential of neuron *i* is denoted by *V*_*i*_. The neurons rest at 0 mV and have a firing threshold (*V*_th_) of 20 mV.

The spike trains generated by neuron *i* is given by 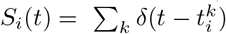 where 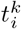 givesthe individual spike times. The transmission delay is denoted by *d*. Individual excitatory postsynaptic potentials have the amplitude *J*_E_ = 0.1 mV, and inhibitory postsynaptic potentials have the amplitude *J*_I_ = − 0.8 mV. The matrix entry *J*_*ij*_ represents the amplitude of a postsynaptic potential induced in neuron i when a spike from neuron j arrives. As multiple synapses can exist from neuron j to neuron i, the amplitude *J*_*ij*_ is an integer multiple of *J*_E_ or *J*_I_, respectively, depending on the type of the presynaptic neuron. ∆*V*_rTMS_ denotes the membrane potential deviation induced by magnetic stimulation which will be introduced in the following section. An action potential is generated when the membrane potential *V*_*i*_(*t*) of the neuron reaches *V*_th_, following which the membrane potential is reset to *V*_reset_ = 10 mV. All parameters are listed in Table 1.

**Table 1.** Parameters of neuron model.

| Parameter | Symbol | Value |
| --- | --- | --- |
| Membrane time constant | $\tau_m$ | 20 ms |
| Resting potential | $V_{\text{rest}}$ | 0 mV |
| Threshold potential | $V_{\text{th}}$ | 20 mV |
| Excitatory postsynaptic potential (EPSP) amplitude | $J_E$ | 0.1 mV |
| Inhibitory postsynaptic potential (IPSP) amplitude | $J_I$ | -0.8 mV |
| Synaptic delay | $d$ | 2 ms |
| Reset potential | $V_r$ | 10 mV |
| Refractory period | $t_{\text{ref}}$ | 2 ms |

### Network model

We modeled an inhibition-dominated recurrent neuronal network (38), with 10000 excitatory and 2500 inhibitory neurons. To study the effects of rTMS on network dynamics and network connectivity, we used both static networks and plastic networks.

### Static network

All inhibitory synapses in the static network have a fixed synaptic amplitude of *J*_*I*_ =− 0.8 mV and excitatory synapses have a fixed amplitude of *J*_*E*_ = 0.1 mV. All synapses among inhibitory neurons, excitatory neurons, and between excitatory and inhibitory neurons are static. These synapses are randomly established with a 10% connection probability. All the neurons in the network receive steady stochastic background input in the form of Poissonian spike trains of *r*_ext_ = 30 kHz. This allows the neurons to have fluctuating subthreshold membrane potential dynamics with pre-determined stable firing rate of 7.8 Hz. The network parameters have been chosen to facilitate an asynchronous-irregular resting state. The network parameters have been listed in Table 2.

**Table 2.** Parameters of network model.

| Parameter | Symbol | Value |
| --- | --- | --- |
| Number of excitatory neurons | $N_E$ | 10000 |
| Number of inhibitory neurons | $N_I$ | 2500 |
| Connection probability | $C_p$ | 10% |
| Rate of external input | $r_{\text{ext}}$ | 30 kHz |

### Plastic network

The plastic network has the same network architecture as the static network, except that the E-E connections were grown from zero following the homeostatic structural plasticity rule implemented in previous works (35, 36, 39). By setting the target firing rate to 7.8 Hz, the network will grow into an equilibrium status driven by the external Poissonian input (*r*_ext_ = 30 kHz), where the average connection probability is around 10% and all neurons fire irregularly and asynchronously around the target rate (7.8 Hz). While using a plastic network, any repetitive magnetic stimulation is only applied after completion of the growth period. Network parameters can be found in Table 2.

### Homeostatic structural plasticity rule

As mentioned above, the connections among excitatory neurons (E-E) followed a homeostatic structural plasticity (HSP) rule, and were subject to continuous remodeling. This rule has been inspired by precursor models by Dammasch (40), van Ooyen & van Pelt (41) and van Ooyen(42). This specific model was previously employed to show cortical reorganisation after stroke (43) and lesion(44), emergent properties of developing neural networks (45) and neurogenesis in adult dentate gyrus (46, 47).

However, we use a more recent implementation of this model in NEST (48) which does not include a distance-dependent kernel, previously used to demonstrate associative properties of homeostatic structural plasticity (35, 39). The authors demonstrated that without the need of an enforced Hebbian plasticity rule, this homeostatic rule can cause network remodeling which displays emergent properties of Hebbian plasticity. Following external stimulation, the affected neurons underwent synaptic remodeling that lead to formation of a cell assembly among these neurons, thus exhibiting activity driven associativity, a distinctive feature of Hebbian plasticity (49). In the present study, we follow this line of thought to propose an alternative mechanism of rTMS induced plasticity.

Each neuron i in this model has a discrete number of dendritic spines (presynaptic elements, 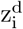) and axonal buotons (postsynaptic elements, 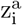), which are paired to form functional synapses. Synapses can only be formed if free synaptic elements are available. Each synapse has a uniform strength of *J*_*E*_ = 0.1 mV. The growth rule we use is a rate-based rule, as implemented in NEST (48). The rule follows the set-point hypothesis, which states that there is a set-point of intracellular calcium concentration that a neuron tries to achieve, in order to maintain stability. Deviations from this set-point level are met by global (whole neuron) efforts to restore it via synaptic turnover. This is in line with experimental results that have shown that neurite growth and deletion are controlled by intracellular calcium concentration (50, 51, 52).

Therefore, in the model of homeostatic structural plasticity used here, the growth and deletion of synaptic elements of a neuron i are governed by its intracellular calcium concentration *ϕ*_*i*_(*t*) = [*Ca*^2+^]*i*. Following each neuronal spike, there is an increase in intracellular calcium concentration by amount, *β*_*Ca*_ through calcium influx. The intracellular calcium concentration decays exponentially with time constant *τ*_*Ca*_ between spikes. The spike train *S*_*i*_(*t*) related intracellular calcium dynamics can be expressed as,

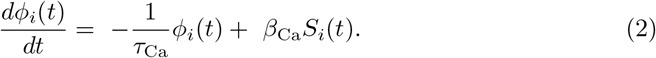

The variable *ϕ*_*i*_(*t*) has been shown to be a good indicator of a neuron’s firing rate (53). According to the synaptic growth rule we use, each neuron i maintains a time-varying estimate of its own firing rate, using its intracellular calcium concentration as a surrogate. This estimate is used by the neuron to control the number of its synaptic elements. When the firing rate falls below the prescribed set-point, indicated by a target firing rate, the neuron grows new synaptic elements to form additional synapses. Following this, freely available pre- and postsynaptic elements are randomly paired with free synaptic elements of other neurons, forming new synapses. These synapses enable the neuron to receive additional excitatory inputs, thus bringing the firing rate back to the set-point. Similarly, when the firing rate rises above the set-point, the neuron breaks existing synapses in order to limit the net excitatory inputs received. The elements from these broken synapses are added to the pool of free synaptic elements. Both the pre- and post-synaptic elements follow this linear growth rule (35, 36),

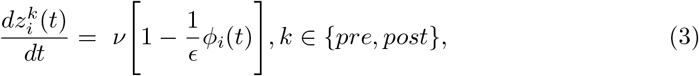

where *i* is the index of the neuron, *ν* is the growth rate and *ϵ* is the target level of calcium. The parameters of the homeostatic structural plasticity rule are listed again in Table 3.

**Table 3.** Parameters of structural plasticity model.

| Parameter | Symbol | Value |
| --- | --- | --- |
| Growth rate | $\nu$ | $0.0039 \text{ s}^{-1}$ |
| Target level of calcium | $\epsilon$ | 0.0078 |
| Time constant for calcium trace | $\tau_{Ca}$ | 10 s |
| Increment on calcium trace per spike | $\beta_{Ca}$ | 0.0001 |

### Model of repetitive Transcranial Magnetic Stimulation (rTMS)

The electrical field induced by rTMS was implemented in the form of current injections into point neurons via a step-current generator in NEST simulator. For mathematical simplification, TMS pulses were modeled as rectangular waves. Each stimulus pulse had a duration of 0.5 ms, modeled after output of conventional rTMS devices, and was depolarizing (monophasic) in nature. Following evidence that rTMS causes changes in spiking behavior of cortical pyramidal neurons (54, 55, 56), we used stimulation intensities that are suprathreshold in nature. This premise allowed us to simplify the role of TMS-induced electrical field in neuronal depolarisation in our simulations. The orientation of the e-field is known to influence the cite of depolarisation in neurons, but since we use spatially simplistic point neurons, the cite of stimulation does not have a specific influence, as long as each stimulus causes an action potential. Previously, a similar approach was taken to model transcranial direct current stimulation (tDCS) (36). Similar to rTMS, there is evidence supporting the therapeutic benefits of tDCS in conditions like major depressive disorder (57, 58) and chronic pain(59, 60). However, unlike rTMS, tDCS is a continuous low intensity stimulation technique, typically not sufficient to cause action potentials. tDCS mainly focuses on modifying the membrane polarity of neurons in order to manipulate their threshold for action potential generation (for an overview, see 61).

The effect of rTMS over networks of neurons has often been described using canonical cortical microcircuit models (62) that include cortical layers II, III, V and inhibitory interneurons. Both inhibitory and excitatory interneurons contribute towards the net effect of TMS, via polysynaptic interactions (63). Accordingly, it has been observed that TMS-induced depolarisation of superficial pyramidal neurons (L2/3) of the canonical microcircuit may lead to recruitment of inhibitory interneurons that project to large pyramidal neurons of layer V (64). Such robust effects of TMS on networks of neurons (54, 65) were observed in our model in the form secondary activation of inhibitory neurons.

In order to investigate the effects of protocol structure, we modeled repetitive stimulation protocols (**Fig 1D**) of different frequencies and intensities. We also modeled the clinically relevant US FDA approved protocol, namely intermittent theta burst stimulation (iTBS) with 600 pulses, described in following sections. Parameters of TMS protocols used throughout this study are summarised in Table 4.

**Fig 1.**
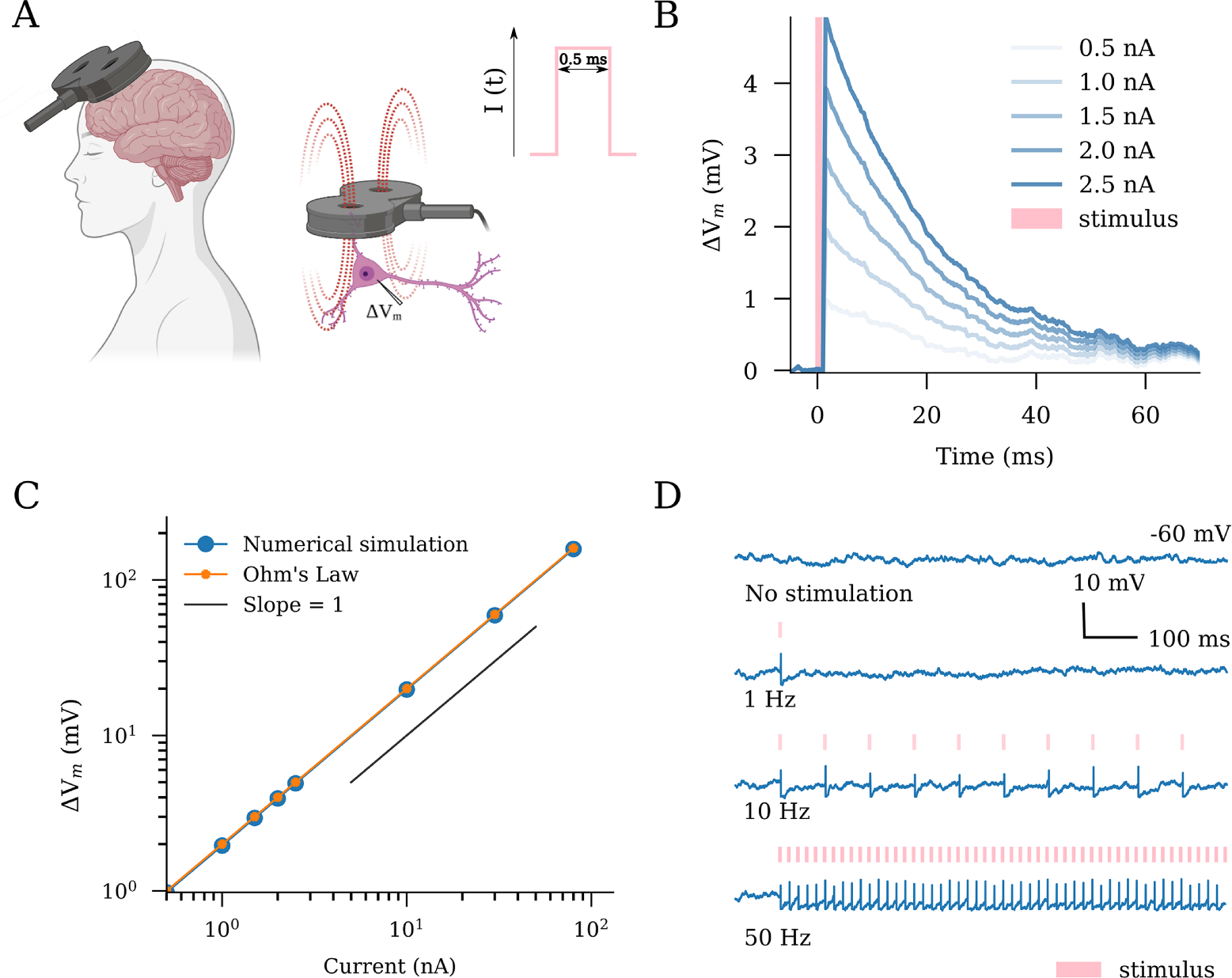
Transcranial magnetic stimulation (TMS) has an immediate effect on the membrane potential dynamics of single neurons. Schematic illustration of TMS in humans and neurons. The TMS-induced electric fields cause depolarization of neurons in the target region. We modeled TMS as rectangular pulse current injections with a duration of 0.5 ms (c.f., standard output parameters of conventional TMS devices). (B,C) Single stimuli produce changes in the membrane potential in a dose-dependent linear manner as predicted by Ohm’s law. (D) Suprathreshold stimulation at different frequencies elicits spiking responses from the stimulated neurons. Created with BioRender.com.

**Table 4.** Parameters of Transcranial Magnetic Stimulation (TMS)

| Figure | Protocol | Frequency (Hz) | $\Delta V$(mV) | Pulse count | E-E synapses |
| --- | --- | --- | --- | --- | --- |
| 1B | single TMS | - | multiple<sup>1</sup> | 1 | - |
| 2C | rTMS | 1, 10, 50 | 0 – 200 | 900 | static |
| 2D | rTMS | 10 | multiple<sup>2</sup> | 100 | static |
| 2E | rTMS | 10, 20, 30, 40, 50 | 20 – 140 | 900 | static |
| 3C | rTMS | 10 | 68 | 900 | plastic |
| 3D | rTMS | 10 | multiple<sup>2</sup> | 900 | plastic |
| 4B | rTMS | 10 | 68 | 900, 3000, 9000, 22500 | plastic |
| 4C | rTMS | 20, 30, 40, 50 | 68 | 300 – 3000 | plastic |
| 4D | rTMS | 5, 10, 15, 20 | multiple<sup>2</sup> | 600 | plastic |
| 5B | rTMS | iTBS | multiple<sup>2</sup> | 600 | plastic |
| 5C | rTMS | iTBS | 68 | multiple<sup>3</sup> | plastic |
| 5D | rTMS | iTBS, cTBS, 10 | 68 | multiple<sup>3</sup> | plastic |

<sup>1</sup> The membrane depolarisation caused are 0.98, 1.96, 2.94, 3.93, 4.92 *mV*.
<sup>2</sup> The membrane depolarisation applied are: $a = 20$ mV, $b = 39$ mV, $c = 68$ mV, $d = 160$ mV.
<sup>3</sup> The pulse numbers used are 300, 600, 900, 1200, 3000 and 9000.

### Numerical experimental protocols

#### rTMS pulse triggering membrane potential deviation

In order to closely observe the response of individual neurons to single rTMS-like stimulus, we modeled single excitatory neurons that receive equal net excitatory and inhibitory Poissonian inputs and therefore maintain subthreshold membrane potential dynamics. Spiking behavior was disabled in the neuron. A single pulse current injection of 0.5 ms duration, which represents a magnetic stimulation pulse in our study, was delivered to the neurons. We observe the membrane potential trace 5 ms before the pulse onset to about 70 ms post the pulse onset. In order to account for randomness and variability, we noted membrane potential traces from 500 individually isolated neurons, all receiving nonidentical but equal net Poisson inputs. The membrane potential traces were averaged to obtain a robust readout. We repeat this experiment for different pulse amplitudes.

#### Theta burst stimulation protocol

Theta burst stimulation delivers bursts of stimuli at a 5 Hz frequency. Each burst consists of three pulses that occur at a 50 Hz frequency. The US FDA approved intermittent theta burst stimulation (iTBS) protocol has a more temporally complex structure. The protocol consists of 600 pulses that last a total duration of 192 s. The pulses are delivered in the theta burst format for 2 s, followed by an 8 s interval. This cycle is repeated 20 times. The continuous theta burst stimulation (cTBS) consists of 600 pulses in the theta burst format delivered in 40 s. The protocol structures can be found in Fig 5A.

#### Analysis and quantification

##### Estimation of membrane potential deviation using Ohm’s Law

The membrane potential deviation in the leaky integrate and fire neurons caused by current injection was estimated using Ohm’s Law. Accordingly, a current pulse of amplitude *A* yields a membrane potential response, *U* (*t*):

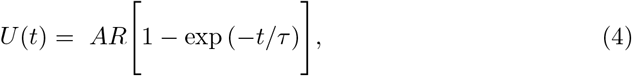

where *R* = 80 MΩ is the membrane leak resistance, *τ* = 20 ms is the membrane time constant of the neuron. In the case of brief pulses, similar to the TMS pulses used in this study, following the current onset, the time course *U* (*t*) of the voltage rises approximately linearly with time:

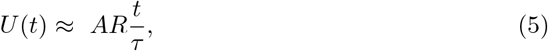

where *t* = 0.5 ms is the duration of the TMS pulse. We used the above formulation to calculate the membrane potential deviation caused by TMS pulses to single neurons.

##### Firing rate

The spiking activity of individual neurons are read out using a spike-detector, as available in NEST. The firing rate is calculated as a spike count average, across defined time-steps of simulation, typically of 1000 ms duration. Mean firing rate of a population are calculated as the arithmetic mean of firing rates of all the neurons from the group.

##### Network connectivity

Connectivity among all or subgroups of excitatory neurons is calculated using an *n × m* connectivity matrix *A*_*ij*_, where n and m represents the total number of presynaptic and postsynaptic neurons, respectively. Each entry in this matrix can either be zero or non-zero positive integers, denoting the total number of synapses from presynaptic neuron j to the postsynaptic neuron i. The connectivity of the whole network or subnetworks were used in the present study for any given time-point t. It is thus calculated as the mean of the whole matrix or corresponding part of it, as follows:

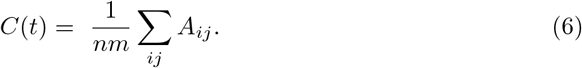

##### Time constant of connectivity saturation

In order to characterise the stimulation duration required to reach connectivity saturation during stimulation, we perform a curve-fitting of the data-points using an exponential function:

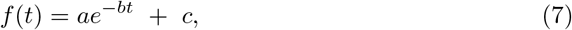

where *τ*_decay_ = 1*/b* represents the time constant of the decay of connectivity during stimulation.

## Results

### Changes in single-neuron membrane potential dynamics and action potential induction in response to transcranial magnetic stimulation (TMS)-like electric stimulation

Multi-scale compartmental modeling demonstrates that the electric fields induced by TMS generally cause changes in the membrane potential of individual principal neurons, eventually resulting in action potential induction and characteristic intracellular calcium level changes (66, 67, 68). Therefore, we first evaluated the effects of TMS-like electric stimulation on the membrane potentials at a single neuron level (Fig 1). For this purpose, single neurons—those receiving balanced excitatory and inhibitory Poissonian spike trains—were stimulated with 0.5 ms rectangular current pulse injections of different amplitudes (Fig 1A and B). A linear interrelation between current injections and membrane potential deviation was observed, consistent with Ohm’s law (Fig 1C). With this approach, implementation of suprathreshold repeated stimulations, i.e., ∆*V* = 68 mV at 1, 10 or 50 Hz, induced robust action potentials in the individual neurons (Fig 1D). We conclude that TMS-like neuronal spiking can be readily induced in our experimental setting.

### Non-linear effects of rTMS intensity on network activity

In realistic applications, TMS activates a network of connected neurons rather than a single neuron. Therefore, we evaluated the effects of increasing stimulation intensities on a subpopulation of neurons embedded in a recurrent network of 10000 excitatory and 2500 inhibitory neurons (Fig 2A). We modeled a focal stimulation that directly affected 10% of the excitatory neurons and studied the network response in terms of the firing rate changes among the following populations: stimulated excitatory neurons (S), non-stimulated excitatory neurons (E), and inhibitory interneurons (I). We first delivered a sample train of rTMS pulses (900 pulses at 10 Hz, c.f., 12, 69, with a pulse intensity that would cause a 68 mV membrane potential deviation) to the subpopulation. As shown in the raster plot, the spiking activity in the stimulated subpopulation was elevated (Fig 2B). We also observed a weaker synchronization throughout the subpopulations during stimulation, indicative of recurrent connectivity. Once stimulation ended, the neurons returned to their baseline Poissonian firing patterns.

**Fig 2.**
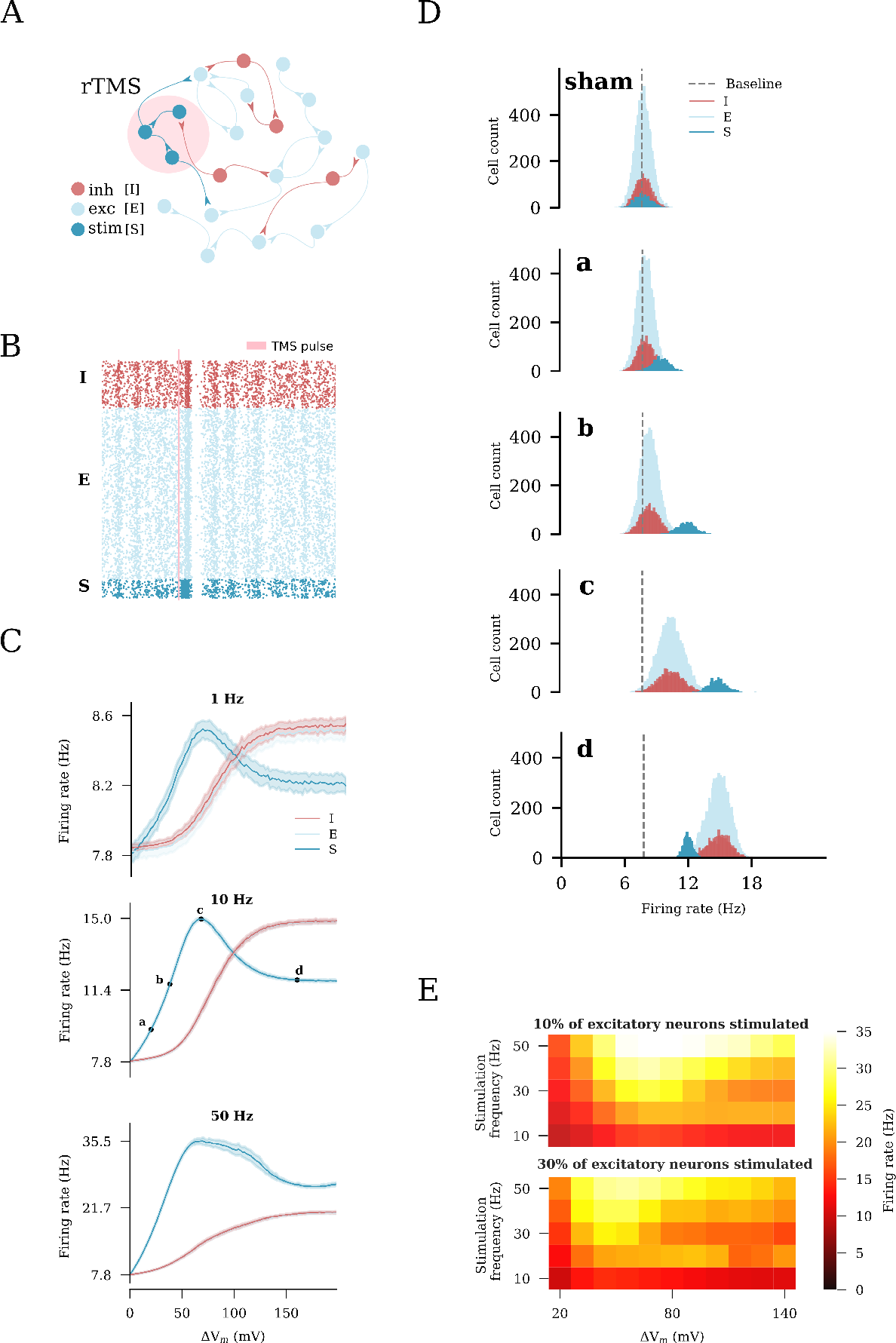
Repetitive transcranial magnetic stimulation (rTMS) changes network activity. (A) Illustration of the recurrent neuronal network with sparsely connected excitatory [E] and inhibitory [I] neurons used in this study. A subset of excitatory neurons [S] is stimulated. (B) rTMS influences the asynchronous, irregular firing state of the stimulated neurons [S], causing them to fire in a synchronous manner. (C) Change in the average firing rate in response to distinct stimulation intensities and frequencies of 10% of excitatory neurons. Four intensities (a: weak, c: peak, d: strong, and b: strong-equivalent) were arbitrarily selected to represent different stimulation intensities. (D) Firing rate histograms for populations E, I, and S at stimulation intensities a, b, c, and d, respectively. (E) Heatmaps summarizing the results of rTMS of 10% (top) and 30% (bottom) of excitatory neurons.

To examine the impact of distinct stimulation protocols on network activity, we performed a series of simulations with varying intensities and frequencies (each at 900 pulses). Examples of the firing rates of the defined subpopulations of interest are shown in Fig 2C. We found that the stimulated population responded at lower stimulation intensities and frequencies (i.e., 1 Hz and 10 Hz), with a proportional increase in the firing rates, which peaked at a stimulus-induced depolarization of 68 mV. With stronger stimulation, the firing rate response of the stimulated subpopulation declined as the firing rate of the inhibitory neurons increased owing to recurrent inhibition. Eventually, a plateau was reached. For higher frequencies (i.e., 50 Hz), changes in the firing rate did not follow the exact same trend as for the lower frequencies (e.g., 1 Hz). This may be attributed to the strong high-frequency stimulation that forced the network to enter into a different stable regime. Nevertheless, the impact of recurrent inhibition on the stimulated neurons was still observable (Fig 2C). The effects of distinct stimulation intensities on the network firing dynamics were carefully examined by plotting the firing rate distributions of the respective sub-populations in response to those intensities (Fig 2D). Weak stimulation did not cause noticeable additional activation of the inhibitory subpopulation. At the peak intensity, the inhibitory neurons were evidently activated. The strong stimulation significantly activated the inhibitory interneurons. The evoked recurrent inhibition had a profound effect on the stimulated subpopulation, resulting in suppression of its firing rate response. The same firing rate of the stimulated neurons was achieved at much lower stimulation intensities that did not recruit inhibition, including strong-equivalent intensity (c.f., Fig 2C). Based on these results, we selected four intensities, characteristic of different states of the network, for further exploration. The resulting values were expressed in terms of the induced changes in the membrane potential of the stimulated neurons and categorized as follows: (a) weak, 20 mV, (c) peak 68 mV, (d) strong, 160 mV and (b) strong-equivalent, 38 mV stimulations.

The results across a wide range of frequencies (10 to 50 Hz) and different stimulation intensities (20 to 140 mV-induced membrane potential change) are summarized in Fig 2E. The described effects on the inhibitory neurons and recurrent inhibition did not depend on the stimulation frequency. We also replicated these results in simulations of a larger subset of excitatory neurons (i.e., when 30% of the principal neurons were stimulated, Fig 2E, bottom). Herein, we observed lower peak firing rates of the stimulated neurons, demonstrating that recurrent inhibition was more effectively recruited when larger populations of neurons were directly stimulated. Taken together, these simulations suggest that an “optimal” stimulation intensity that effectively increases the firing rate of stimulated neurons exists. Exceeding this intensity leads to further recruitment of inhibition, which dampens the activity of the stimulated excitatory neurons. Lower strong-equivalent stimulation intensities can be determined at which the same effects on the firing rates of stimulated neurons are observed, without major effects on network inhibition.

### Structural remodeling of network connectivity in response to rTMS

We switched to a plastic network that remodels its connections in an activity-dependent homeostatic manner (Fig 3). This network follows a plasticity rule where an increase in the firing rate of excitatory neurons leads to retraction and disconnection, while a reduction in the firing rate promotes outgrowth and formation of new excitatory contacts between principal neurons (Fig 3A; c.f., 35, 36). In this study, stimulation was performed after an initial growth stage, which allowed the network to reach a steady state with 10% connectivity between the excitatory neurons and a mean firing rate of 7.8 Hz (Fig 3B). We applied a 10 Hz stimulation protocol consisting of 900 pulses at peak intensity to a subset of 10% of excitatory neurons (c.f., Fig 2B). As described above, the stimulation elicited an instant increase in the firing rates of the stimulated neurons as well as non-stimulated excitatory and inhibitory neurons (Fig 3C). This sudden increase in the firing rates was accompanied with a homeostatic structural response where the principal neurons reduced existing input synapses to restore baseline activity. This disconnection was most prominently observed among the stimulated neurons, but also occurred between the stimulated and non-stimulated excitatory neurons (Fig 3C). The end of stimulation, which was also marked by a sudden drop in the net input received by the non-stimulated excitatory and inhibitory neurons, led to an instant drop in firing rates. This was followed by the formation of new connections that compensated for the now reduced network activity. As activity returned to baseline, a reorganization of network connectivity became evident: The stimulated neurons showed significantly more connections among each other (S-S), while the connection between the stimulated and non-stimulated neurons (S-E) was reduced; The connectivity among the non-stimulated neurons (E-E) remained unaltered. These simulations suggest, that rTMS-like electric stimulation can have distinct effects on the connectivity among and between stimulated and non-stimulated neurons, as reported before (c.f. Fig 2 of 36).

**Fig 3.**
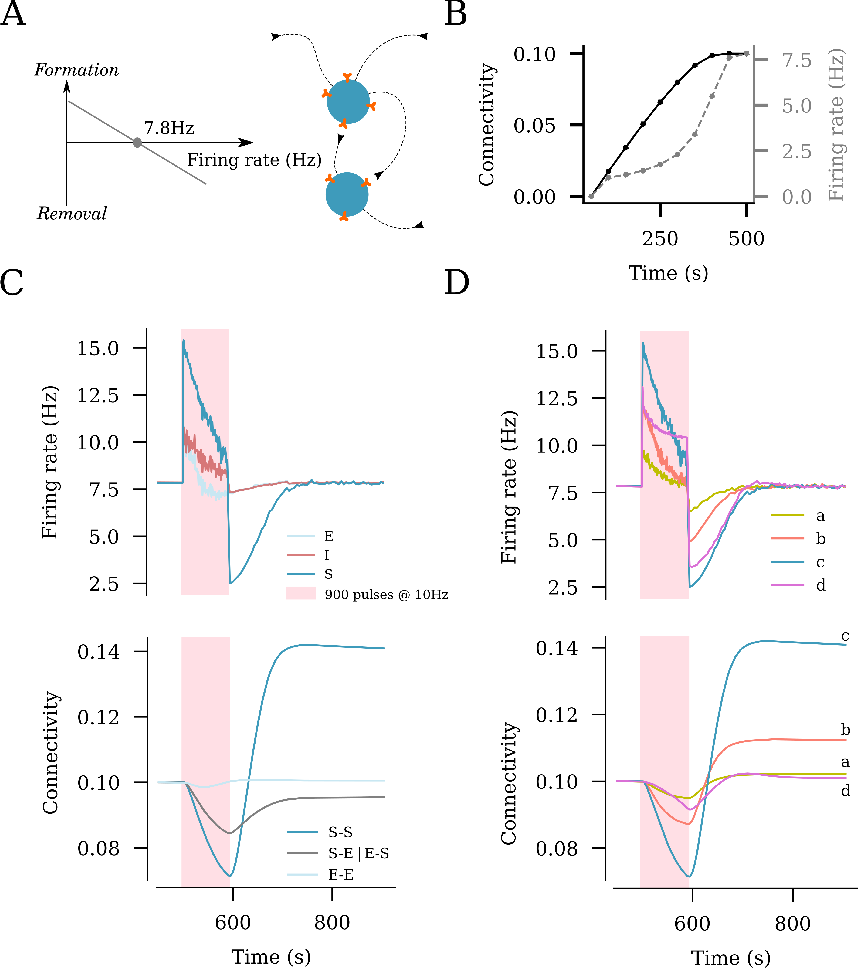
rTMS induces structural remodeling of stimulated networks. (A) Homeostatic structural plasticity assumes negative feedback of neuronal activity on its connectivity with other neurons: A high firing rate removes synapses between excitatory neurons, and a low firing rate promotes synapse formation. (B) Poissonian input stabilizes the firing rate and connection probability prior to stimulation. (C) Effects of a 10 Hz stimulation protocol consisting of 900 pulses on the firing rate and structural remodeling [i.e., connectivity between stimulated neurons (S–S), between non-stimulated excitatory neurons (E–E), and between stimulated and non-stimulated neurons (S–E and E–S)]. (D) Effects of the same stimulation protocol on the firing rate of stimulated neurons and connectivity between stimulated neurons at the four representative amplitudes from Fig 2C [i.e., weak (a), strong-equivalent (b), peak (c), and strong (d)].

**Fig 4.**
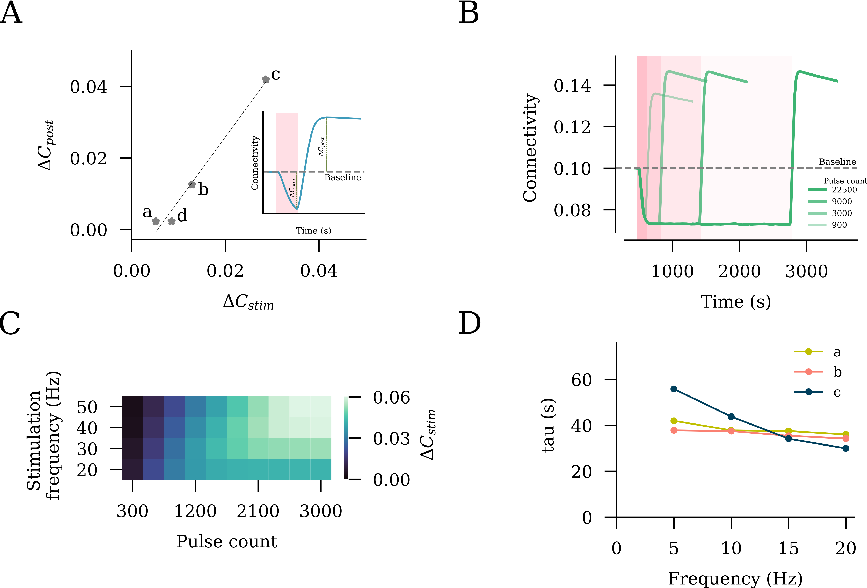
rTMS intensity and pulse number affect the structural remodeling of stimulated networks. (A) Interrelation between the connectivity drop during stimulation (∆*C*_*stim*_) and connectivity increase post stimulation (∆*C*_*post*_). (B) Stimulation outcomes from different pulse numbers of 10 Hz stimulation at peak stimulation intensity (c, as defined in Fig 2C). (C) Saturation points, expressed as the total pulse numbers required, are summarized for a range of frequencies. (D) Time constants of connectivity decay (*τ*_decay_) were extracted by fitting an exponential function to stimulation connectivity drop among stimulated neurons (S–S).

**Fig 5.**
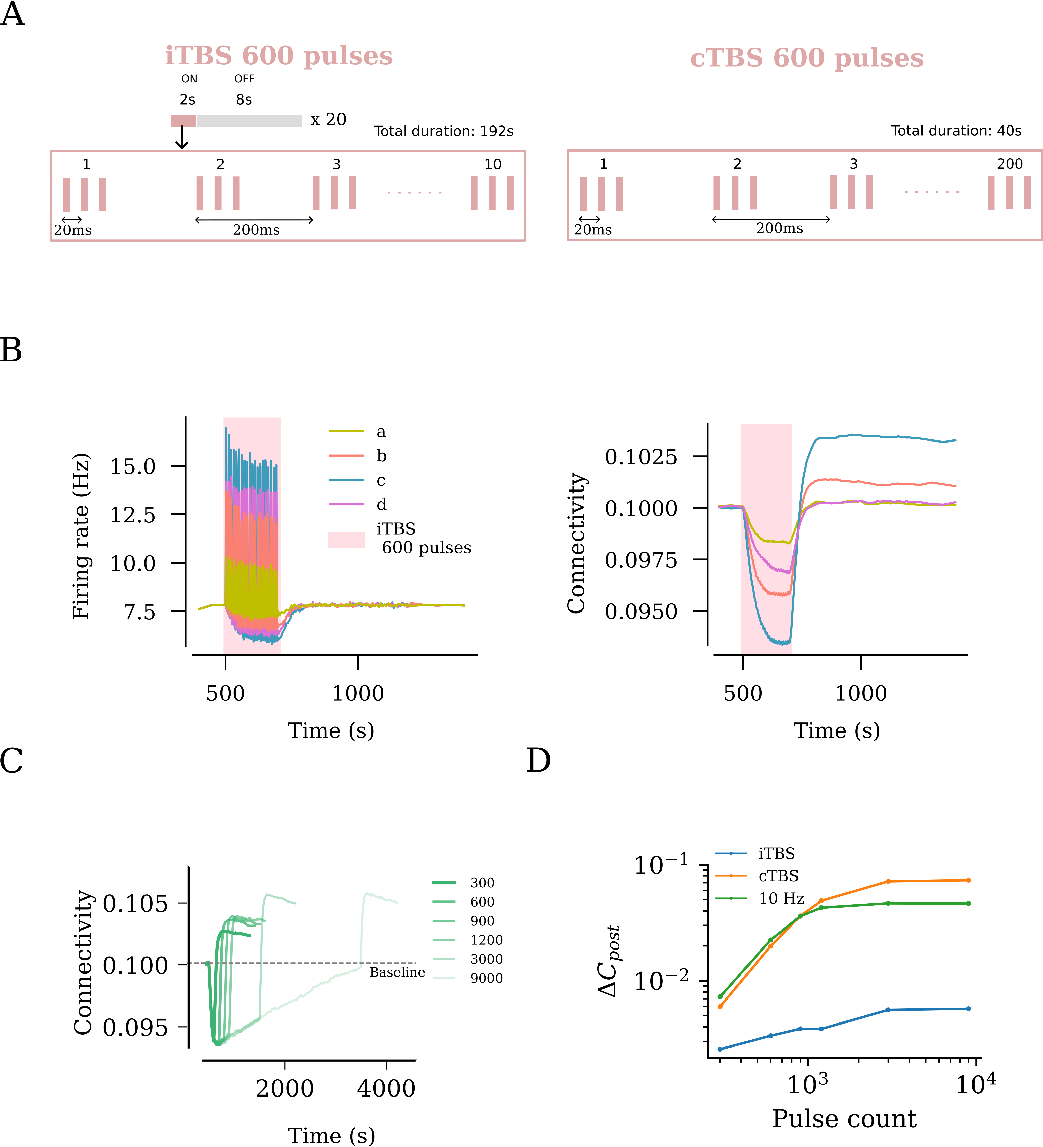
rTMS leads to duration and intensity dependant overstimulation for intermittent Theta Burst Stimulation (iTBS). US FDA approved iTBS protocol consists of 600 pulses distributed across ON times of 2 s and OFF times of 8 s. The ON times consist of ten bursts of stimulus pulses at 5 Hz, where each burst consists of 3 pulses occurring at 50 Hz. (B) iTBS applied at peak amplitude (c, as defined in Fig 2C) resulted in the strongest firing rate response and the largest network connectivity upshoot. (C) iTBS at increasing stimulation duration (i.e., pulse numbers) was found to cause increasing values of post-stimulation connectivity upshoot among stimulated neurons. This trend was tested for iTBS, cTBS and 10 Hz and is summarised as log-log plots in (D).

### Dose-dependent effects of rTMS on structural network remodeling

We also assessed the outcome of the distinct stimulation intensities on homeostatic structural plasticity and network connectivity (Fig 3D). The same stimulation protocol (10 Hz, 900 pulses) was applied with weak, peak, strong, and strong-equivalent intensities (c.f., Fig 2B and C). As shown in Fig 3D, the largest change in the connectivity among the stimulated neurons was seen in response to the peak amplitude (i.e., a 68 mV membrane potential increase). The weak amplitude elicited a small response in neural activity, and only minor changes in lasting connectivity were observed (Fig 3D). The strong and strong-equivalent amplitudes yielded different effects on connectivity. The network receiving strong-amplitude stimulation failed to rapidly restore its activity to baseline by homeostatic structural plasticity during stimulation, which was reflected in a weaker overall connectivity change. This may be attributed to the recurrent inhibition recruited by a strong electric stimulation, which then affected the stimulated neurons. This phenomenon was not observed in the strong-equivalent stimulation, while a considerable remodeling of network connectivity was noted (Fig 3D).

### Influence of the stimulation duration on network remodeling

We noted that the extent of network connectivity changes after stimulation depended on the degree of reorganization caused during stimulation. Indeed, a proportional interrelation was observed between these two parameters (Fig 4A). This observation had important implications for the stimulation duration, including the number of pulses applied at a given frequency. The finding suggests that once the increase in the firing rate is compensated, the application of additional pulses will not have a further effect on the outcome of intervention, at least not in terms of lasting changes in network connectivity after stimulation.

To explore this hypothesis, we applied 10 Hz stimulations of different durations to 10% of the excitatory neurons and assessed the trajectories of connectivity among the stimulated neurons (Fig 4B). We observed an increasing post-stimulation peak connectivity with an increasing stimulation duration. However, this relationship did not hold beyond a certain point. For 10 Hz stimulation, we found that stimulation beyond *∼*3000 pulses did not contribute to further changes in the peak connectivity. This allowed us to conclude that the connectivity change has reached a saturation point, and 10 Hz stimulation for longer durations would not have a stronger effect on network connectivity (Fig 4B). Indeed, the outcome of a stimulation with 22500 pulses was comparable to that observed with 3000 and 9000 pulses, as shown in Fig 4B.

We followed up on this observation by extending our simulations to include a range of frequencies from 10 Hz to 50 Hz, as summarized in Fig 4C. The trend of connectivity saturation was maintained, with lower frequencies taking larger pulse numbers to reach the saturation point. Considering that the pulse number is equal to the total stimulation duration multiplied by the frequency, it is therefore unknown what the role of stimulation duration is. We thus extracted the time constant of decay (*τ*_decay_) by fitting exponential curves to connectivity data obtained during stimulation. The *τ*_decay_ values across different frequencies at a fixed stimulation intensity were comparable, with a trend of inverse proportionality in case of peak stimulation intensity. We deduce that the total stimulation duration has a major impact on the net stimulation outcome, irrespective of the frequency.

### Effects of the clinically approved iTBS protocol on network activity and connectivity

Finally, we evaluated the effects of the clinically approved iTBS protocol, which we found to have a more complex stimulation pattern with inter-train intervals (Fig 5A). We systematically applied the four relevant stimulation intensities, namely weak, peak, strong, and strong-equivalent, and assessed the changes in network connectivity (Fig 5B). Similar to what we observed with 10 Hz stimulation, the weak and peak stimulation intensities led to small and large changes in connectivity, respectively. Comparatively, the strong-equivalent intensity induced intermediate changes in connectivity, while the strong stimulation intensity led to only small changes in connectivity.

We then evaluated distinct stimulation durations, including the pulse numbers at peak stimulation intensity, and found that a plateau was reached between 600 and 1200 pulses, with 900 pulses showing approximately the same effect as 1200 pulses on network connectivity (Fig 5C). An additional increase in connectivity was evident at 1500 pulses, indicating that unlike the 10 Hz stimulation protocol, the iTBS protocol may assert additional effects when large numbers of pulses are applied. Indeed, the simulations with 3000 and 9000 pulses (c.f., Fig 4B) confirmed this suggestion (Fig 5D). Notably, the effects of the iTBS protocol on structural remodeling were weaker than those of the pulse-matched 10 Hz stimulation protocol (Fig 5D). This difference may be attributed to the inter-train interval of the iTBS protocol. Consistent with this suggestion, pulse-matched continuous TBS (cTBS) induced structural remodeling that exceeded the effects of iTBS and 10 Hz stimulation (Fig 5D). Taken together, these results emphasize the relevance of proper selection of stimulation parameters, specifically the stimulation intensity and pulse number, where “overdosing” may have negative or at least no additional desired effects.

## Discussion

In this study, we explored the effects of rTMS on network dynamics and connectivity using simulations of an inhibition-dominated recurrent neural network with homeostatic structural plasticity. rTMS was found to increase the activity of neurons and induce characteristic changes in network connectivity. These effects of rTMS depended on the stimulation intensity, frequency, and duration. Differential effects of rTMS were observed in the stimulated and non-stimulated neurons; the connectivity among the stimulated neurons increased, while disconnection between the stimulated and non-stimulated neurons was observed. Our simulations suggest that recurrent inhibition, which is recruited at high stimulation intensities, may counter rTMS-induced neural activation and plasticity. We also observed that increasing the number of stimulation pulses beyond a certain point may saturate the structural network reorganization. Thus, optimal stimulation protocols where no additional desired effects will be observed by further increasing the intensity of stimulation or number of TMS pulses may exist.

However, for the FDA-approved iTBS protocol, we observed an additive effect on the changes in network activity at larger pulse numbers. We attribute this effect to the complex pattern of the iTBS protocol, specifically the inter-train intervals. iTBS at 900 pulses seems to be more effective than iTBS at 600 pulses in our simulations. Notably, however, the effects of iTBS on the structural remodeling of the stimulated networks were weaker than those of pulse-matched 10 Hz stimulation or cTBS. Taken together, our results suggest a new mechanism of rTMS-induced plasticity that does not depend on LTP-like plasticity and synaptic weight changes. This rTMS-induced homeostatic structural plasticity is sensitive to specific parameters of the stimulation protocol, emphasizing the need for a careful standardization and a systematic experimental assessment of dose–response relationships in rTMS-based basic and clinical studies.

Although direct experimental evidence on the human neocortex is still lacking, it seems well established in the field that rTMS changes cortical excitability by modulating excitatory and inhibitory neurotransmissions (70, 71, 27). However, the effects of rTMS on cortical excitability—measured as changes in the amplitudes of motor evoked potentials—return to baseline within 90 min after stimulation. Therefore, it is unlikely that rTMS-induced LTP or long-term-depression (LTD) is the major or sole mechanism underlying the therapeutic effects of rTMS that can last weeks or months after stimulation (72, 73). Yet, clinical protocol designs are often based on studies reporting rTMS-induced LTP- or LTD-like plasticity (74, 75, 7). Herein, we used computational modeling to explore an alternative biomechanism of rTMS that is based on homeostatic plasticity and structural remodeling of neuronal networks-homeostatic structural plasticity. Homeostatic plasticity involves activity-dependent negative-feedback mechanisms that aim at maintaining neuronal networks within a stable operational range (76, 77, 78): An increase in network activity leads to weakening of excitatory synapses, strengthening of inhibitory synapses, and therefore shifting in the excitability of neurons. Previously, Gallinaro and Rotter demonstrated emergent associative properties of homeostatic structural plasticity, via activity-driven formation of neuronal ensembles (35). Consistent with these previous findings and with the use of a similar computational approach, the present results suggest that rTMS triggers an activity-dependent disconnection of neurons that enables the formation of new excitatory synapses and leads to a profound structural remodeling of stimulated networks.

While some experimental evidence supports the existence of homeostatic structural plasticity (79, 80,81, for overview, see 82), its biological significance and the underlying molecular mechanisms warrant further investigation. In our previous work, in which we used live cell microscopy to study the effects of rTMS on dendritic spines of cultured hippocampal CA1 neurons, we did not find any significant changes in the synapse numbers, including spine density changes following 10 Hz repetitive magnetic stimulation (12). This is consistent with the finding of a recent *in vivo* two-photon imaging study demonstrating subtle structural changes in dendritic spines in response to repeated sessions of low-intensity rTMS (14). Synaptic (un)-silencing could be one of the biological implementations of homeostatic structural plasticity (82, 83, 84).

Synapses that are typically found on small dendritic spines or filopodia containing mainly NMDA receptors are referred to as “silent”, as NMDA receptors are blocked by magnesium ions at resting membrane potential. They can be activated after the accumulation of depolarizing AMPA receptors (85, 86, 87, 88). Indeed, our previous work revealed that 10 Hz repetitive magnetic stimulation promotes the accumulation of AMPA receptors at small preexisting spine synapses and triggers the growth of these presumably silent dendritic spines (12). Thus, rTMS may mediate homeostatic structural plasticity by conveying to neurons the ability to remove or form functional synaptic connections by regulating the accumulation of AMPA-receptors at preexisting synapses, without the need to recruit the complete molecular machinery to remove or form new spines and/or synapses.

In a network without structural plasticity, we observed a non-linear relationship between the stimulation intensity and neuronal firing rate changes. This non-linearity in the firing rate response can be attributed to recurrent inhibition. We observed increasing feedback inhibition in response to higher stimulation intensities. This effect had a major impact on the outcome of rTMS-induced structural plasticity. Accordingly, we defined four critical stimulation intensities for closer examination: weak, peak, strong and strong-equivalent. At amplitudes below the peak value, the inhibitory subpopulation was not strongly activated. Meanwhile, with stimulation stronger than the peak amplitude, stronger recurrent activity recruited the inhibitory subpopulation, which consequently inhibited the stimulated subpopulation, causing a weaker firing rate response. Indeed, stimulation stronger than the peak amplitudes yielded weaker effects on structural remodeling than did stimulation at a lower intensity, despite their comparable effects on the firing rates of the stimulated neurons. In general, this highlights the important role of inhibitory networks in rTMS-induced plasticity.

Experimental evidence suggests that single pulse TMS inhibits neocortical dendrites by directly activating axons within the upper cortical layers, which leads to the activation of dendrite-targeting inhibitory neurons in the neocortex of mice (89). Moreover, our previous work showed that 10 Hz rTMS remodels inhibitory synapses: Dendritic but not somatic inhibition as well as the strength, sizes, and numbers of inhibitory synapses were reduced after stimulation (16). These findings emphasize that rTMS also induces structural changes in inhibitory networks. In line with these findings, rTMS has been shown to trigger the remodeling of visual cortical maps (90, 91). However, the direct effect of stimulation on inhibitory neurons and homeostatic structural plasticity of inhibitory synapses remain elusive. The dose-dependent effects on specific inhibitory neuron types and their impact on rTMS-induced structural remodeling of excitatory and inhibitory synapses warrant further investigation (92, 93, 94). Regardless of these considerations our findings suggest that strong stimulation may lead to less effective structural remodeling of stimulated networks as compared with weak stimulation that causes equivalent changes in the firing rates.

Our model also makes predictions relevant for translational applications of rTMS. Based on our findings, we propose a model of “connectivity saturation”. Stimulating networks of neurons initiates homeostatic synaptic remodeling that leads to loss in connectivity among the neurons. The end of stimulation period is followed by further synaptic remodeling causing increase in connectivity among the affected neurons. We used an exponential function to fit the trajectory of connectivity during the stimulation period and extracted time constants of connectivity decay, *τ*_decay_. This value can be roughly interpreted as the least time required to attain structural equilibrium during stimulation. This translates to the maximum remodeling that is attainable once stimulation is turned off. We found that the *τ*_decay_ values were comparable for low stimulation intensities across a wide range of frequencies, emphasizing the relevance of the stimulation duration rather than the pulse numbers. At the peak stimulation intensity, we found a slight frequency dependency indicating, that lower frequencies take a longer time to achieve connectivity saturation. A similar connectivity saturation was not observed in the iTBS protocol. However, the effects of iTBS on structural remodeling were much weaker than those of pulse matched 10 Hz stimulation or cTBS. This effect may be attributed to the inter-train intervals, which enabled the network to rewire during the stimulation protocol. Translational frameworks that combine computational models and *in vitro* and *in vivo* animal studies with experiments in healthy individuals are required to confirm and extend the relevant predictions on dose-response interrelations obtained in our computer simulations. However, computational models may already help in advising protocol designs, which are currently mainly based on studies reporting rTMS-induced LTP- (or LTD-) like plasticity.

## Acknowledgments

We thank Dr. Júlia V. Gallinaro for valuable discussions. We thank Dr. Sandra Diaz-Pier from the Forschungszentrum Jülich for support on features of NEST. We thank the Freiburg University Computing Center for computational resources and support that contributed to the results of this study. This work was supported by NIH (1RO1NS109498 to AV) and by the Federal Ministry of Education and Research, Germany (BMBF, 01GQ2205A to AV).

